# Robust and Remote Measurement of Heart Rate Based on A Surveillance Camera

**DOI:** 10.1101/2021.05.13.443967

**Authors:** Yiming Yang, Chao Lian, Hongyu Zhang, Liangjiang Li, Yuliang Zhao

## Abstract

Arrhythmia is a marked symptom of many cardiovascular diseases (CVDs). The accurate and timely detection of heart rate can greatly reduce the harm of arrhythmia to people. However, it is still a challenge to robustly and remotely measure heart rate in daily life due to the changing environmental conditions during measurement, such as the varying light intensity, the movement of people, and the uncertain distance between the sensor to people. In this study, we propose a method to accurately measure human heart rate within a distance of 4.5 meters under different light intensities by simply using a surveillance camera. After a 20-second color video of a person’s hand is captured by the camera, a method based on the Fast Fourier Transform (FFT) algorithm is designed to extract the blood volume pulse wave to calculate the heart rate. According to the comparison between the real heart rate and results measured by electrocardiography (ECG), the proposed method achieves an accuracy of 95.8% when the measurement is performed within a distance of 4.5 meters and 90% when within 5.0 meters. Our experiments show that when the illuminance varies between 100-1000 lux (lighting level indoor), we still get the correct results. Our experiments also demonstrate that the proposed method accurately obtain heart rate even when the light intensity is below 32 lux (300-500 lux in a workplace environment). The method’s strong adaptability to changing environmental conditions makes it applicable to many scenarios, such as homes of the elderly, classrooms, and other public spaces.

## I. INTRODUCTION

Cardiovascular diseases (CVDs) are a leading cause of death today [1]–[3]. Around 17.9 million people died from CVDs in 2016, representing 31% of all global deaths [4]. A large number of clinical studies have shown that irregular heart rate is a complication of CVDs (such as coronary heart disease [5], hypertension [6] and heart failure [7]), and can be used to predict the occurrence of CVDs [8], [9].

Frequent heart rate monitoring has a very positive effect on the prevention of CVDs. The traditional heart rate measurement method, electrocardiography (ECG), requires patients to stick electrodes on the body or wear a chest strap. ECG will cause some irritation and discomfort to patients. Moreover, the device is not that easy to carry, which is inconvenient for most usage scenarios, such as homes or offices.

Currently, quite a number of remote non-contact heart rate measurement methods have been developed. Examples include methods based on microwave Doppler radar [10], fast hyperspectral imager with 25 spectral channels in the NIR [11], and thermal imaging [12], but they all require using expensive and specialized hardware.

Remote photoplethysmography (rPPG) is a non-invasive, remote, low-cost measurement method. It uses a camera to collect the reflected light from the skin and analyzes it to obtain heart rate. Existing rPPG methods use Independent Component Analysis (ICA) [13], Principal Component Analysis (PCA) [12], Butterworth pass band filter [14], Random Forest Regression (RFR) [15], and Fast Fourier Transform (FFT) [11], [16] for heart rate measurement. These methods cannot always effectively detach the heart rate signals from the noises brought by the complicated environment and subjects. Therefore, their illumination robustness and the available distance are very limited. The parameters of these methods are compared in Tab. 1.

**TABLE I.** Comparison of different rPPG methods.

| Core algorithm | Shooting time<br>/second | Research on<br>illumination<br>robustness<br>Y/N | Research on<br>illumination range<br>Y/N | Measuring range<br>/meter | Study |
| --- | --- | --- | --- | --- | --- |
| ICA | 60 | N | N | 0.5 | [13] |
| PCA | 30 | N | N | 1 | [12] |
| Butterworth pass<br>band filter | 60 | N | N | <3 | [14] |
| RFR | 35 | N | N | 1 | [15] |
| FFT | 30 | N | N | 0.8-1 | [11] |
| FFT | 30 | N | N | 1-2 | [16] |
| Filtering+ FFT | 20 | Y | Y | >4.5 | This work |

From the above comparison, we clearly see two main problems of these methods. First, detection of heart rate under poor lighting conditions and drastically changing illuminance levels is under-researched. Second, their effective measurement distance is shorter than the normal maximum room width (3.1 m)[17], which causes much inconvenience to the detection of heart rate. *Sebastian Zaunseder et al*. [18] conducted a comprehensive evaluation and analysis of a large number of iPPG technologies. Most of the studies were carried out in a relatively ideal environment, and there has been a lack of research on heart rate monitoring and measurement uncertainty [19], [20] in a complex environment. Therefore, our study has taken into consideration different illuminances, distances, human body conditions, and other factors.

*TP Sacramento* [21] and *Dangdang Shao* [22] *et al*. studied the feasibility of using a fixed color channel as raw data. We have found a simple method for quick filtering based on the use of fixed color channels. In this study, we propose a method that uses a home surveillance camera for heart rate detection. It can detect heart rate within a distance of 4.5 meters from the patient, is adaptable to low-light environments and has excellent illumination robustness. Our method is superior to the other rPPG methods in three special operations. First, we found that the image processing operation by R channel minus G channel can eliminate most environmental noises. This makes our method sufficiently robust against illumination and insusceptible to drastic illuminance changes. This point is supported by our heart rate detection experiments. Second, identifying the strongest signal position on the skin and setting it as the ROI, we greatly improved the measurement accuracy and extended the measurement distance. This fully validates the ability of our method to eliminate noises in the case of illumination and distance changes. Third, by using the above two algorithms, we used the FFT algorithm to filter a lot of environmental noises and calculate the heart rate. Finally, we used the proposed method to measure the heart rate of the ROI at several different distances and illuminance levels.

By reducing the no-signal area and enhancing the heart rate signal, this method can successfully extend the measurement distance. When the user appears in the shooting range of the camera, the system can detect the heart rate based on subtle changes in the user’s skin color, and upload the data to the cloud for analysis. This can be a great reminder to patients suffering from CVDs to seek medical diagnosis as early as possible. With the ability to remotely monitor users’ heart rate, the proposed method will be helpful in maintaining human health and can be used in many different scenarios to determine the possibility of a user suffering from CVDs. Moreover, it can be reasonably expected that the use of a powerful camera will make the proposed method much better able to help CVD patients in a variety of areas, such as at-home health care, personal fitness, electronic commerce, and financial trading.

## II. METHOD

### A. EXPERIMENTAL PROTOCOL

The contraction and relaxation of the heart causes periodic changes in arterial blood flow, which causes changes in skin color (Fig. 1). Therefore, heart rate can be measured by analyzing the subtle differences in skin color. A large number of experiments have proved the effectiveness of this method. This research also uses a camera to collect images of the skin and measures the heart rate by analyzing these time-series images.

**FIGURE 1.**
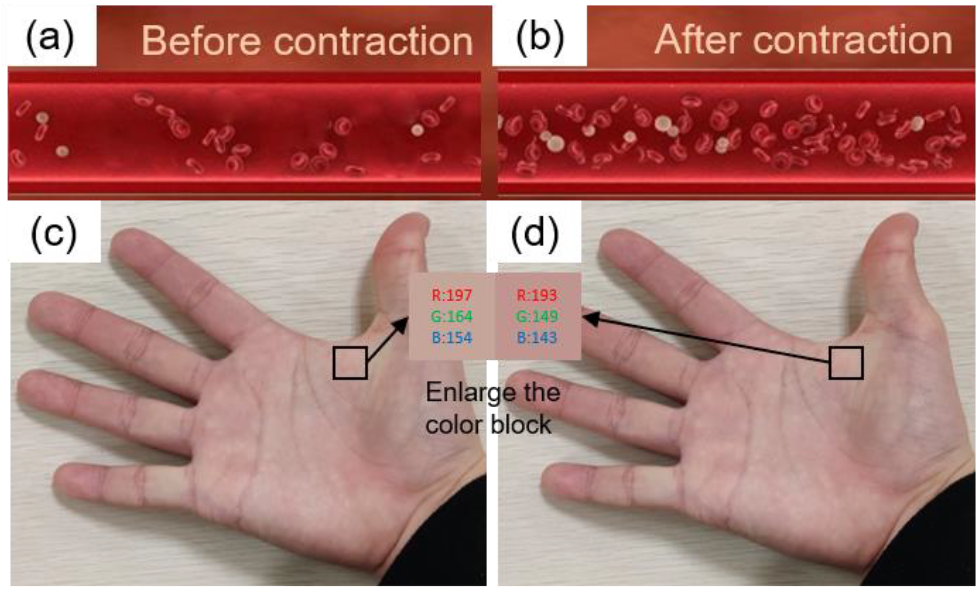
Schematic of the blood vessels before (a) and after (b) heart contract. The color of the hand before (c) and after (d) heart contract.

The detection device used in this study consists of a 3-lead electrocardiogram (Texas Instruments’ ADS1292) and an ordinary camera (Sony IMX345 sensor, F/2.4, 52 mm, 1/3.6, 1 μm, AF). The camera worked at a frame rate of 60 frames/s. All videos were recorded in color space with a resolution of 1920 × 1080. During the measurement, the subject’s hand was placed flat under the camera. The light intensity was measured by HABOTEST’s HT620L. In total, five healthy volunteer subjects (three males and two females at the age of 47 with a yellow skin color) without any known CVD participated in the experiment. Informed consent was obtained from each subject. The vertical distance between the camera and the hand was less than 6.4 meters. To verify the effectiveness of the proposed method, a 3-lead ECG was used to measure the subjects’ heart rate simultaneously.

The subjects stuck the electrodes of the ECG signal collector to the body part as shown in Fig. 2 (ECG signal used as the benchmark). The subjects put their hand flat facing the camera. At the beginning of the experiment, the ECG signal collector and the camera were started at the same time to ensure time synchronization. The ECG signal collected by the ECG signal collector and the video captured by the camera were transferred to the computer for further analysis.

**FIGURE 2.**
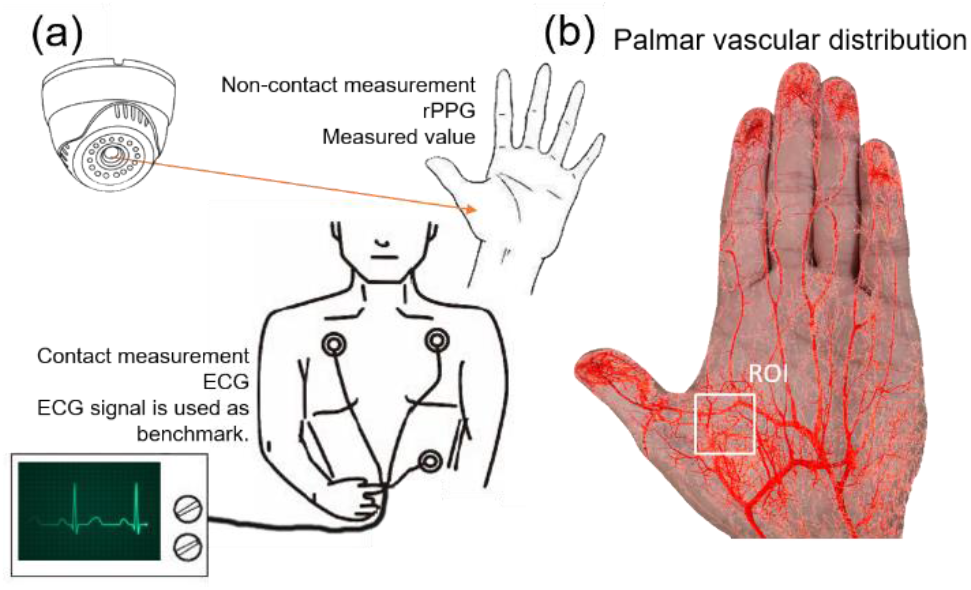
(a) Experimental setup. (b) Blood vessel distribution in the palm.

### B. NOISE FILTERING

The video of the hand was shot and then preprocessed into a sequence of images. First, each frame of the video was saved as an image in matrix form. Second, after the palm was recognized from the image, the ROI was cut from the palm image as shown in Fig. 3. Although Fig. 3A and Fig. 3B are the two images with the largest color difference, it is still difficult to distinguish them only with the naked eyes. The ROI-selection was performed automatically by a computer program developed based on the Media Pipe multimedia machine learning model. As shown in Fig. 3, 24 automatic feature points can be detected, and the hand ROI can be tracked and selected automatically. During the whole process, there was no need for the palm to stay still.

**FIGURE 3.**
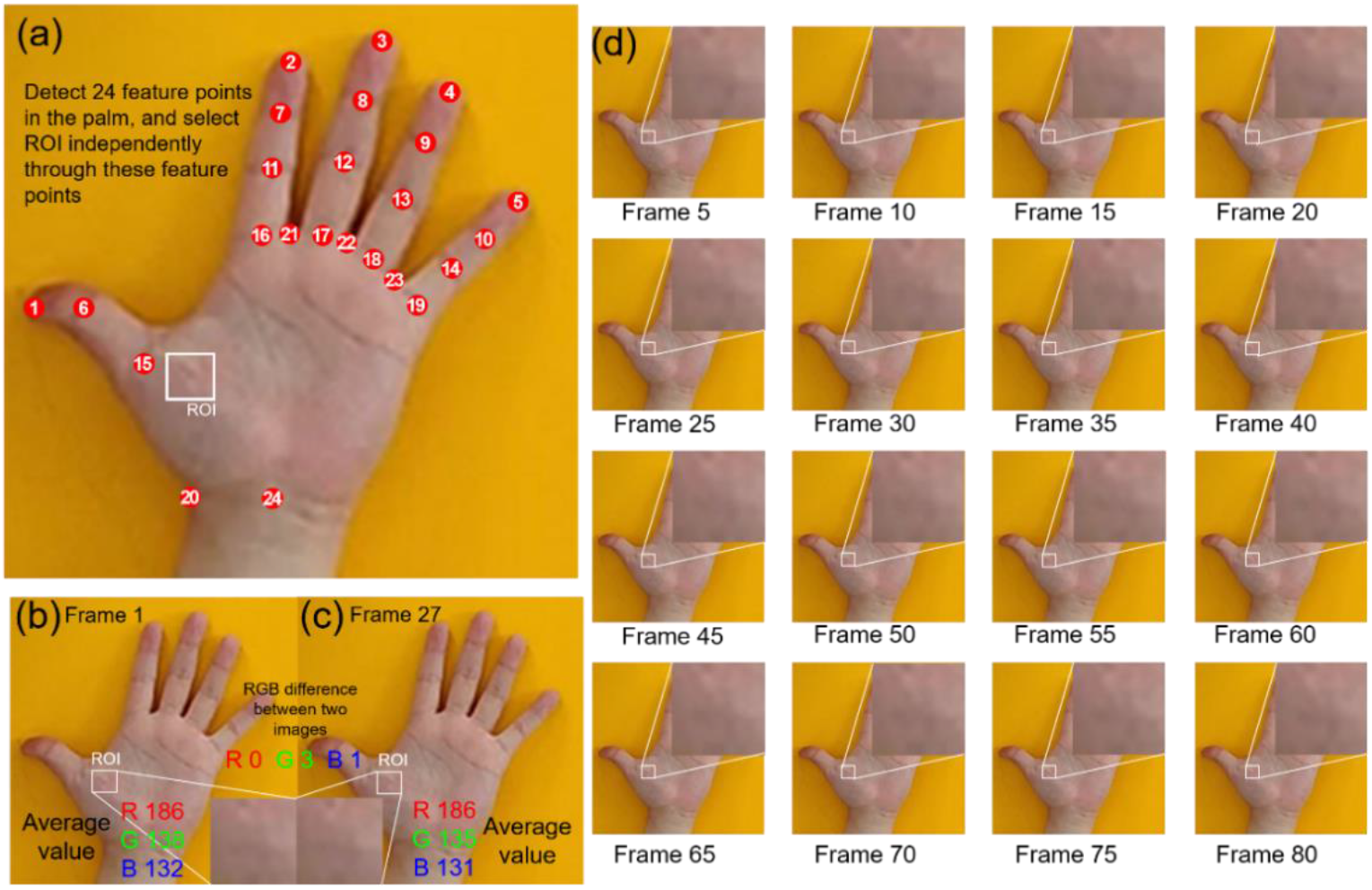
(a). 24 feature points identified on the palm. (b, c) Comparison images between two frames. (d) The captured sequence images. The video was shot at 60 fps.

A main limitation with this type of rPPG measurement is the environmental noises, including the white noise from the camara, unbalanced illumination, and differences in the transparency of the subjects’ skin. Therefore, techniques such as ICA and PCA [15]–[17], [23], [24] have been frequently used to filter the original data. These algorithms can work properly in many scenarios, but their measurement accuracy will drop significantly when the illuminance changes drastically. To minimize the influence of illumination changes on data, a method based on the absorption of different lights by blood was proposed to reduce noise.

Once the video had been preprocessed, the *R*(*k*), *G*(*k*), and *B*(*k*) values were obtained. Specifically, *k* represents the number of frames of the images, *R*(*k*) the sum values of the ROI in the red channel, *G*(*k*) the sum values of the ROI in the green channel, and *B*(*k*) the sum values of ROI in the blue channel.

When the environmental light changes, the RGB value changes as well. A sharp change in the environmental light will cause the noise to be much larger than the heart rate signal, thereby leading to a failure to obtain the heart rate. Therefore, many studies [12]–[16] directly use RGB as data and cannot cope with sudden changes in contrast. Studies have shown that blood can hardly absorb any red light [23], [24]. This means that changes in the red value is an indicator of environmental light changes [25]–[27]. We proposed a filtering method by calculating the difference between R and G values to eliminate the sharp change in RGB from changes in the environmental light. The sharp change in RGB was eliminated by subtracting the G value from the R value. The filtered signal is expressed as

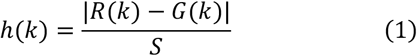

Where *h*(*k*) is the filtered signal, and *S* is the area of the measurement area. The experimental results showed that this formula works better than all the other formulas, as illustrated in Fig. 4. Following the above operations, the video was converted into the signal *h*(*k*), which shows the best SNR. The following operations were performed based on the signal *h*(*k*).

**FIGURE 4.**
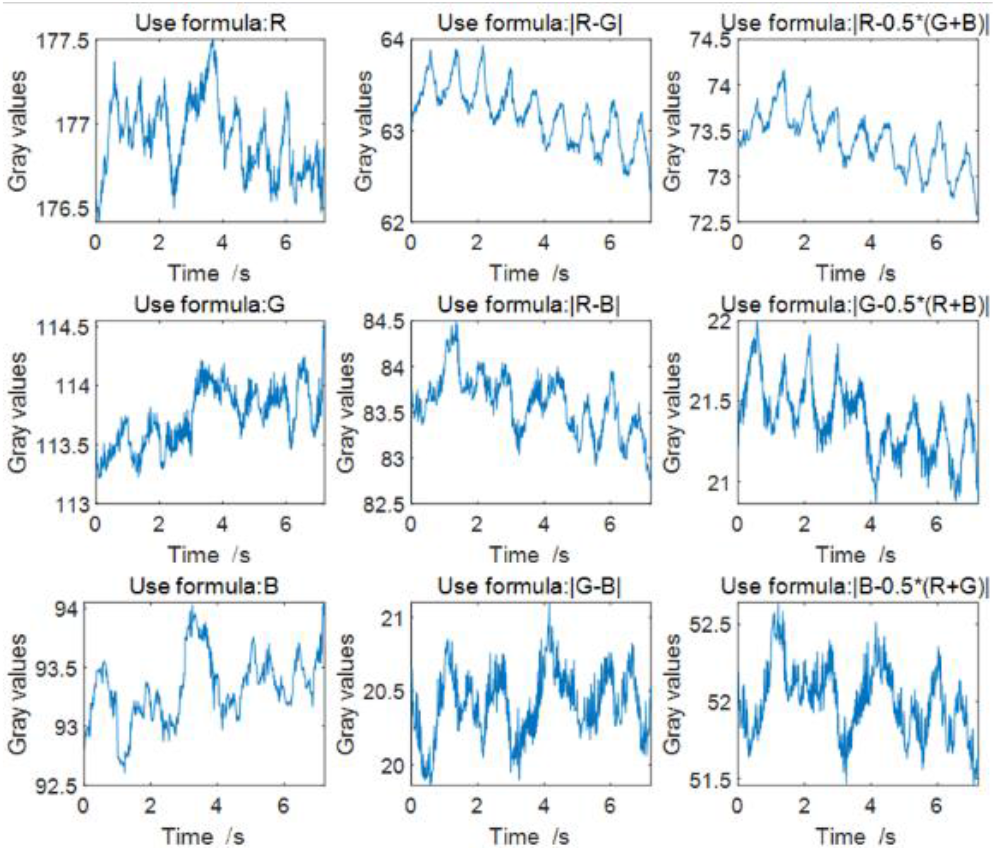
Gray value change curves obtained using different formulas.

### C. FFT FOR HEART RATE MEASUREMENT

Heart rate from the filtered data *h*_(*k*)_ is obtained after filtering the noise by subtracting the G value from the R value. After the early R-G processing, the noise was greatly suppressed. Then, FFT was used to obtain the heart rate quickly and accurately. The FFT formulas are as follows:

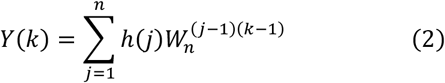

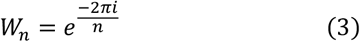

Where *n* is resolution of FFT, and *Y*(*k*) is a vector sequence obtained after the FFT calculation. The modulo operation was performed on *Y*(*k*) to obtain *M*(*k*) (the frequency domain data of the filtered signal). The filtered signal in frequency domain is expressed as *M*(*k*) = |*Y*(*k*)|.

Following the above operations, the filtered signal *h*_(*k*)_ (time domain signal) was converted into *M*(*k*) (frequency domain signal). *M*(*k*) shows the amount of signal (Bd) in each given frequency band within a frequency range. The physiological range of human heart rate is between 40 and 220 beats per minute (bpm) [28]. Therefore, the *Hr_L_* value was set to 40 bpm and *Hr_H_* to 220 bpm. Then, heart rate values above *Hr_H_* and below *Hr_L_* were deleted as noise from the measured frequency domain signal. The frequency domain signal *M(k)* at this time became *M*(*k*) (*Hr_L_* < *k* < *Hr_H_*). The peak of the FFT corresponds to the largest amplitude in the time domain. The non-periodic noise will not show any peak in the frequency domain. However, the heart rate is periodic, and a peak can be found from the frequency domain. This means that the measured signal can be transformed into a peak in the spectrogram, that is, the heartbeat cycle. The maximum value of *M*(*k*) between *Hr_L_* and *Hr_H_* was taken as the heart rate measured by rPPG.

### D. SEARCH OF ROI

A suitable ROI must contain enough blood flow signal to obtain accurate heart rate. There are two main methods for searching the ROI. One is to set the entire skin as the ROI [13]. This method reduces the calculation burden of the device, but weakens the overall SNR with large low signal area. The other one is to select the area with intensive blood vessels on the skin as the ROI [15], [24]. In this study, we investigated the SNR maps of the palms of five subjects of different genders and occupations as shown in Fig. 5.

**FIGURE 5.**
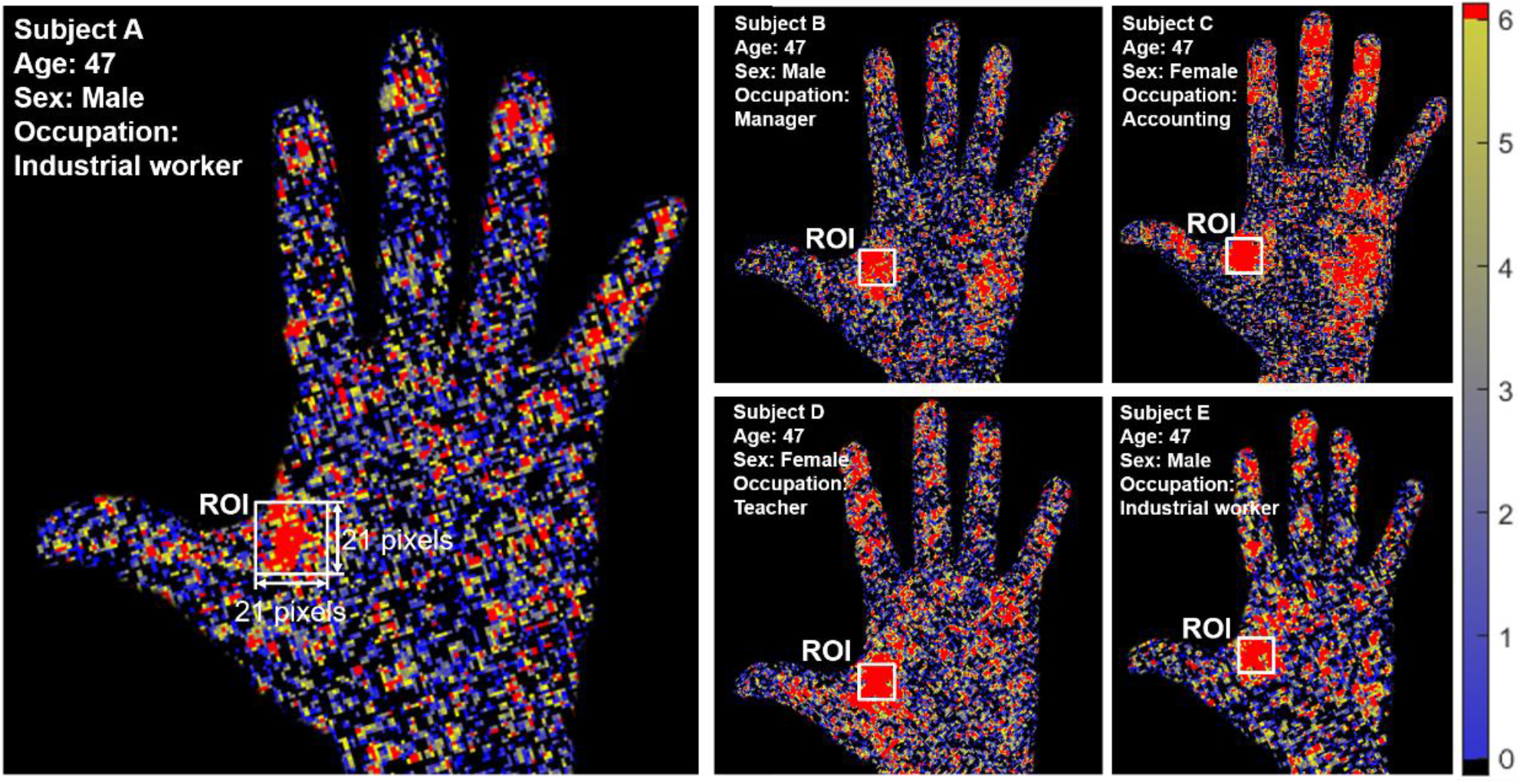
SNR maps of the palms of five subjects of different genders and occupations.

First, Eq. 4 was used to collect the filtered data *h*(*k*) of this pixel. Where *k* represents the number of frames of the images, *R*(*k*) is the intensity values in the red channel, and *G*(*k*) is the intensity values in the green channel.

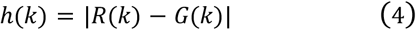

Eq. 2, Eq. 3, and *M*(*k*) = |*Y*(*k*)| were used to perform the FFT operation on the signal of this pixel to obtain the content of the signal at frequency domain *M*(*k*).

Third, Eq. 5 was used to calculate the average value of the frequency domain from *Hr_L_* to *Hr_H_*.

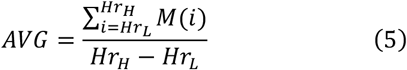

Fourth, ECG was used to obtain the true heart rate *Hr_T_* of the subject as control, and Eq. 6 was used to calculate the SNR.

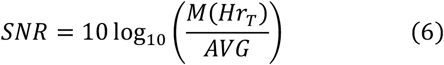

As shown in Fig. 5, different colors indicate pixels with different SNRs. The black color indicates pixels with an SNR lower than 0, and the red color indicates those with an SNR higher than 6. Pixels with the highest SNR occurred most frequently in the area marked by a white square. Therefore, this area was selected as the ROI. This area accounted for about 4~6% of the whole palm.

## III. RESULTS AND DISCUSSION

### A. RESULTS

Using the above-mentioned methods, we analyzed the hand video shown in Fig. 3, which was shot at a distance of 3.1 meters (The normal maximum room [17]) at an illuminance of 150 lux (Homes illumination [29]). The results of the operations based on our method are illustrated in Fig. 6. The waveforms in Fig. 6A show the changes in the average intensity of pixels in the ROI within 10 consecutive heartbeat cycles. However, the result could not show the periodicity of the heart rate clearly. Comparatively, the signal filtered by subtracting the G value from the R value describes the periodicity of the heart rate clearly in Fig. 6B. Comparing with the ECG data in Fig. 6C, we clearly see that the periodicity of our results coincides with that of the ECG results. To verify this, the signal amplitudes of the ECG and rPPG results at different frequencies were calculated, and then compared using the FFT algorithm as shown in Fig. 6D. The coincidental peaks of the two waveforms (69 bpm) obtained by the two methods demonstrated that the heart rate measured by our rPPG method agrees well with that by ECG. We also applied our proposed method to the forehead with an illuminance of 150 lux and at a distance of 3.1 m. We found that the measurement accuracy was reduced slightly from 99.99% to 98.87%, which was mainly because the signal-to-noise ratio was reduced from 4.88 to 2.53.

**FIGURE 6.**
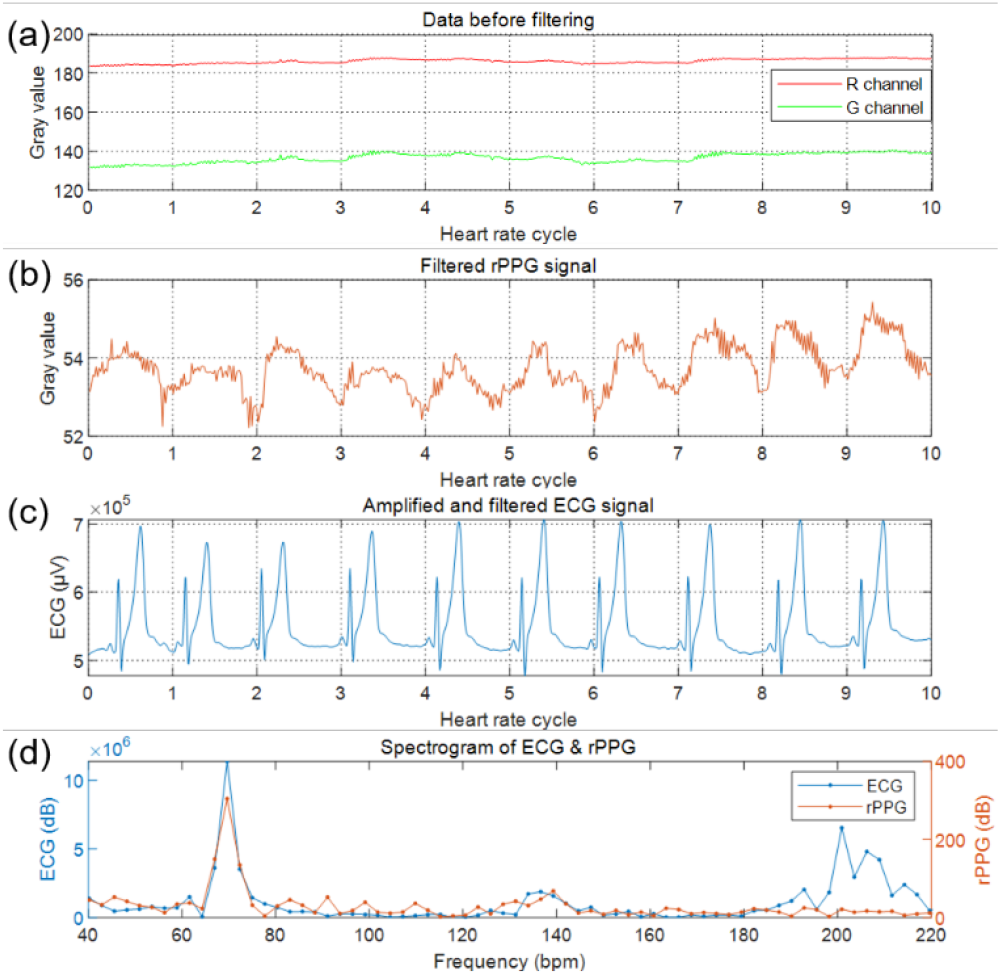
Waveforms of the data measured by rPPG before (a) and after (b) filtering. (c) Waveforms of the data measured by ECG after filtering. (d) Amplitude distribution of waves of different frequencies obtained by analyzing the data measured by rPPG and ECG using FFT.

**FIGURE 7.**
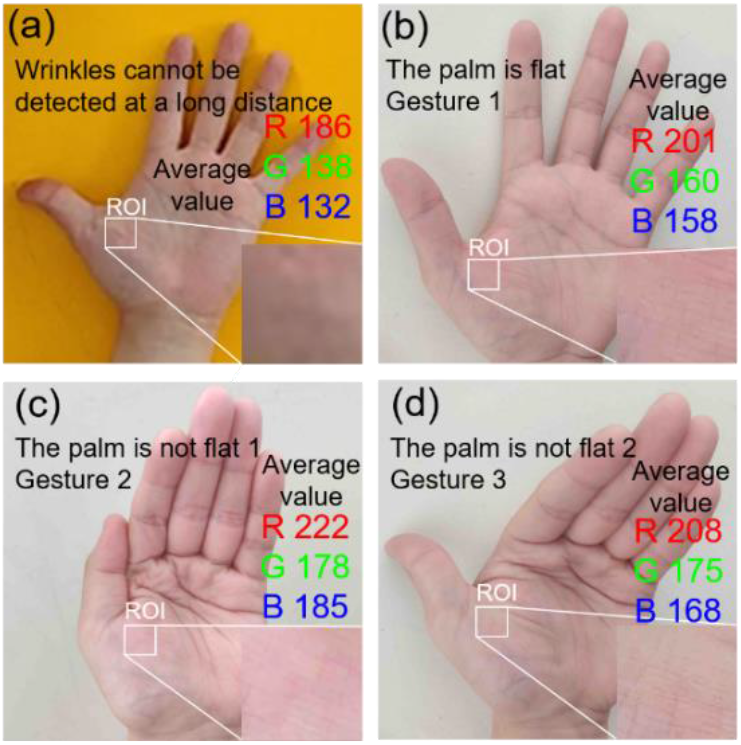
(a) The camera cannot detect the folds. (b-d) Different measuring gestures.

### B. MEASUREMENT ACCURACY WITH DIFFERENT GESTURES

To explore the influence of palm folds on the experimental results, 3 sets of different gesture data (the specific gestures are shown in Supplemental Fig. 7 (a), (b) and (c)) were tested. When the distance exceeds 1.2 meters, the camera we used cannot capture enough details of the wrinkles (Supplemental Fig. 7 (a)). Therefore, when the distance exceeds 1.2 meters, the folds will not affect the results. We set the experiment with distances of 50 cm, 100 cm, 120 cm, 200 cm, 400 cm and 450 cm, and an illuminance of 150 lux (Homes illumination [24]). Accuracy comparison of the different gestures at these distances demonstrated that the folds of the palm will not affect the results (Tab. 2).

**TABLE II.** Accuracy rates of different gestures at different distances.

| Distance /cm | Gesture 1 | Gesture 2 | Gesture 3 |
| --- | --- | --- | --- |
| 50 | 99.99% | 99.99% | 99.99% |
| 100 | 99.99% | 99.99% | 99.99% |
| 120 | 99.99% | 99.99% | 99.99% |
| 200 | 99.99% | 99.99% | 99.99% |
| 400 | 99.39% | 99.89% | 99.99% |
| 450 | 97.60% | 97.87% | 98.20% |

Comparison of palms with no wrinkles (gesture 1) and folds (gesture 2 and 3) demonstrated that the presence of wrinkles will not affect the results of the experiment.

### C. MEASUREMENT WHEN HEART RATE IS ABNORMAL

To verify the validity of our method in measuring abnormal heart rates, a controlled experiment was set up between exercising and resting. We randomly selected 20 people as experimental samples, and set the measurement illumination to 150 lux (Homes illumination [24]). Heart rates were measured during relaxation, after 20 deep squats, and after 40 deep squats at a distance of 400 cm. The experimental results are shown in Tab. 3.

**TABLE III.**
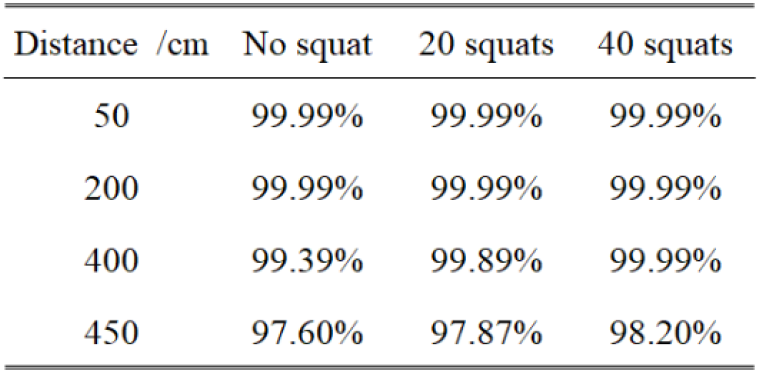
The influence of different levels of exercise on measurement accuracy.

The validity was 99.39% for no squat group, 99.89% for the 20 squats group, and 99.99% for the 40 squats group. With the increase of the amount of motion, the measurement validity increased. Because after strenuous exercise, the blood flows faster in the blood vessels, which leads to more valid heart rate measurements after deep squats [30], [31].

### D. MEASEREMENT ACCURACY WITH DIFFERENT ILLUMINANCE LEVELS

To explore the influence of illuminance on the experimental results, seven more experiments were performed under different lighting conditions at a distance of 3.1 meters (The normal maximum room [17]) from the same subject. The illuminance was kept in the range of 283.5-284.3 lux (normal office lighting [29], [32]), 100.3-101.4 lux (dark overcast day[29], [33]), 52.4-53.5 lux (living room lighting [32]), 32.4-32.5 lux, 14.8-16.8 lux (dark surroundings), 9.8-10.8 lux (twilight), and 3.4-3.5 lux for each of the seven experiments.

Fig. 8 shows that the measurement accuracy increased exponentially with the increase of illuminance and reached 96.71% when the illuminance increased to 32.4 lux. When the illuminance increased to 100.03 lux, the measurement accuracy increased slowly and reached a plateau at ~ 99.99%. We use the standard deviation as the error bar in Fig. 8.

**FIGURE 8.**
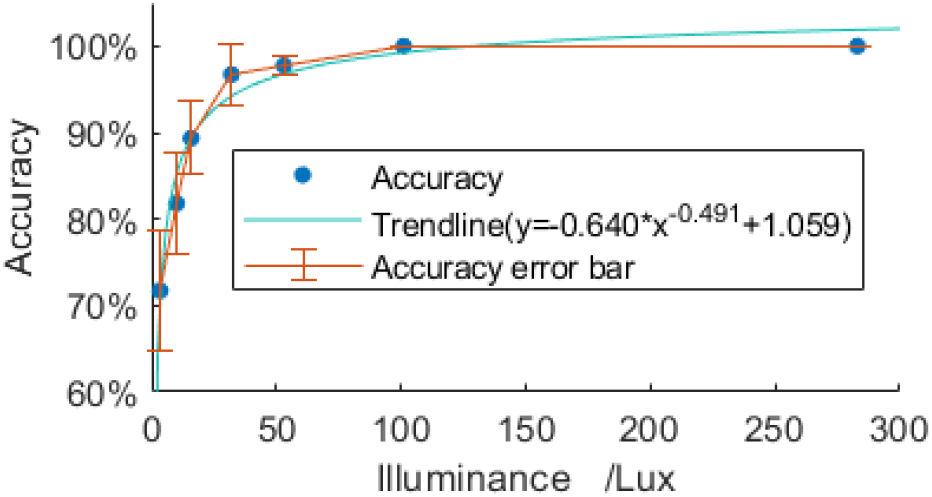
The influence of illuminance on signal recognition. (The experiment of each illuminance level was carried out 10 times)

**TABLE IV.** The influence of different illuminance on measurement accuracy.

| Ambient light intensity /Lux | Accuracy | Standard deviation of accuracy |
| --- | --- | --- |
| 283.5~284.3 | 99.99% | 0.01 |
| 100.3~101.4 | 99.99% | 0.01 |
| 52.4~53.5 | 97.76% | 1.18 |
| 32.4~32.5 | 96.71% | 3.54 |
| 14.8~16.8 | 89.35% | 4.21 |
| 9.8~10.8 | 81.82% | 5.77 |
| 3.4~3.5 | 71.72% | 6.88 |

### E. ILLUMINATION ROBUSTNESS VERIFICATION

To evaluate the illumination robustness and reliability of our rPPG method, we performed a series of experiments by introducing two types of light noises with drastically increased and decreased illuminance at a distance of 3.1 meters (The normal maximum room [17]) from the same subject. So, the two types of light noises are as follows: 1) illuminance decreased drastically (from 1000 lux to 100 lux), and 2) illuminance increased drastically (from 100 lux to 1000 lux). Fig. 9 shows the gray values of R and G channels before and after data filtering. The signal before filtering changed drastically with illuminance changed. No periodic changes were observed from the signal before filtering. With filtered signal, periodic changes is clearly observed, and its changes affected by changes in illuminance were very small. This experiment proves the robustness of our method.

**FIGURE 9.**
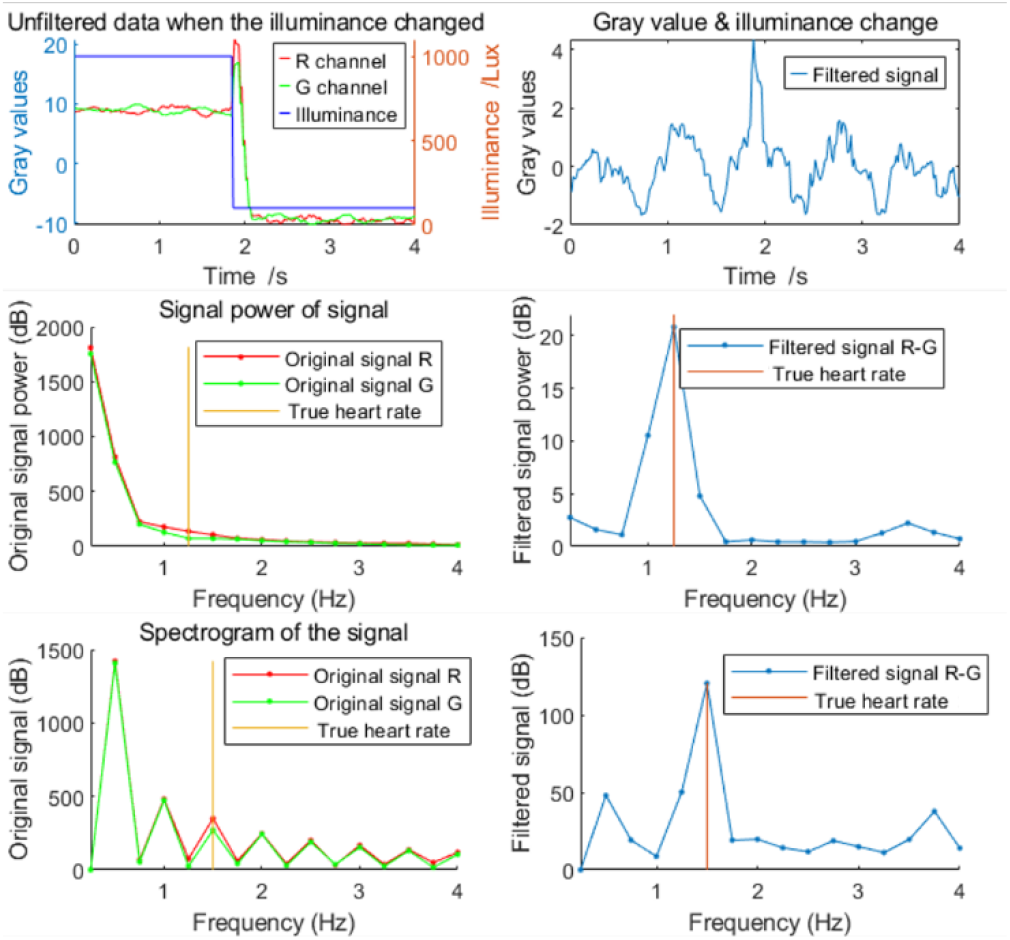
Data obtained when the illuminance changed.

**TABLE V.**
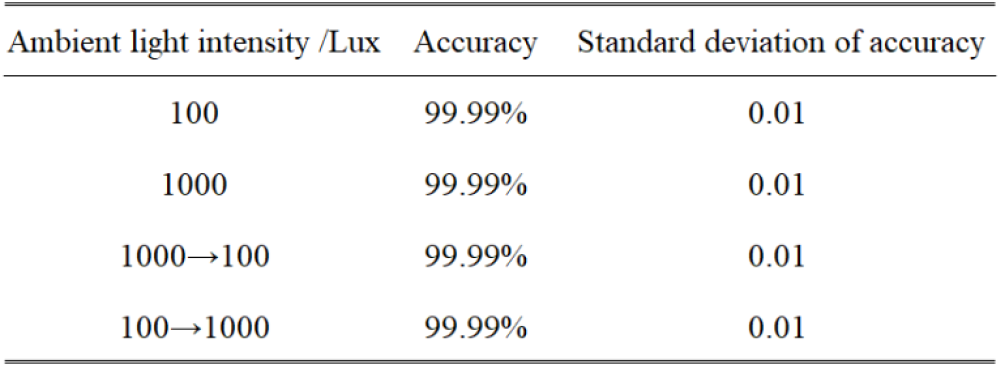
The influence of the illuminance changes on measurement accuracy.

### F. MEASUREMENT ACCURACY WITH DIFFERENT DISTANCES

We performed a series of experiments to test the measuring accuracy with different distance between the camera and the subject. The same environmental light (150 lux, homes illumination [29]) was used for the 12 groups, and the measurement distances were set to 50 cm, 100 cm, 120 cm, 200 cm, 250 cm, 300 cm, 350 cm, 400 cm, 450 cm, 500 cm, 550 cm, and 600 cm, respectively.

Fig. 10 shows that the measurement accuracy decreased with the measurement distance. When the measuring distance within 350 cm, the accuracy was kept above 99%. When the distance increased to 450 cm, the accuracy decreased slowly but was still at a high level (above 95%). However, when the distance was over 550 cm, the camera experienced decreases in its sensitivity and resolution. In this case, the measurement accuracy dropped to 57.6%. We use the standard deviation as the error bar in Fig. 10.

**FIGURE 10.**
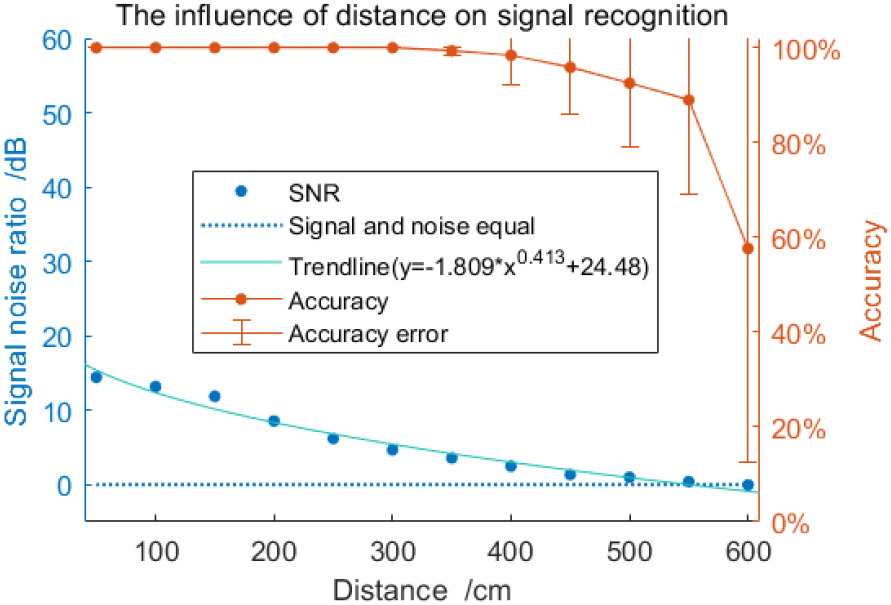
The influence of measurement distance on signal recognition and SNR (30 volunteers participated in the measurement at each distance).

**TABLE VI.**
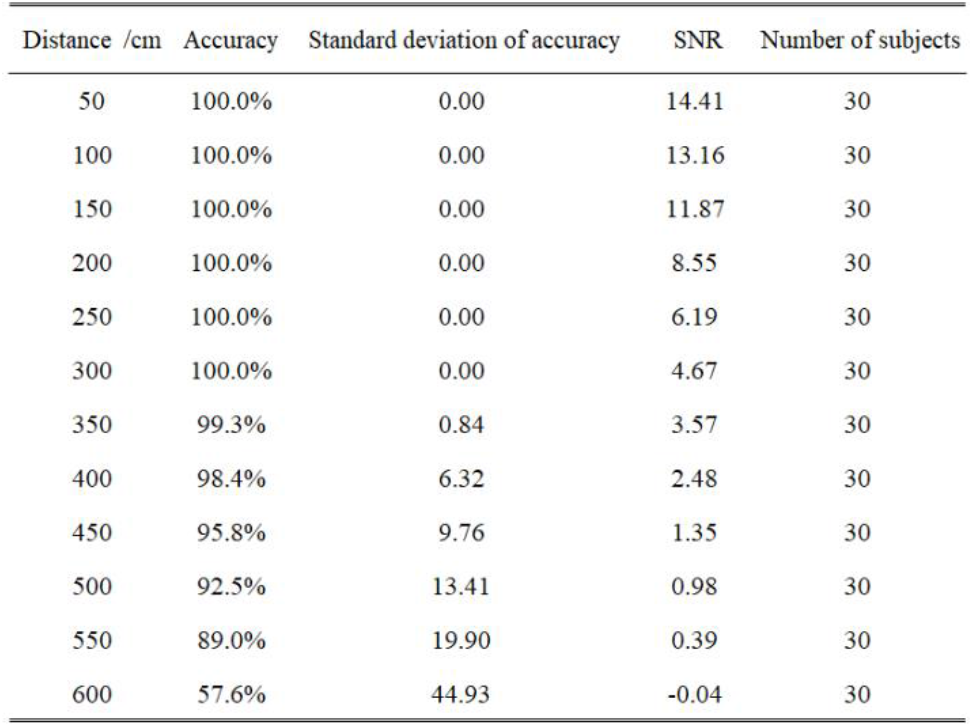
The influence of different distances on measurement accuracy.

## IV. FUTURE WORKS

In our future works, we will use surveillance cameras with infrared capability for data collection, and shift our focus to near-infrared light filtering and data analysis. We will work on multi-person heart rate measurement on remote monitors in public places. We will optimize the filtering algorithm to enable longer-distance measurement, and introduce face tracking for multi-activity target ROI selection. We will collaborate with hospitals to collect data from a large number of patients and realize the prediction of some common CVDs through deep learning.

## V. CONCLUSION

We found that many remote heart rate measurement methods require a stringent lighting environment and the measurement to be performed at a distance. We used an R channel minus G channel method to detect the relative value of the gray value change in the ROI area, and used FFT to obtain the heart rate. The combination of these two methods greatly reduce the influence of illumination changes on the measurement accuracy. We used the above method to calculate the SNR of each pixel in the video, and identified the area with the largest SNR as the ROI. This greatly reduced the need for illuminance and allowed for extended measurement distances.

We measured the heart rate at a distance of 3.1 meters under stable and sufficient lighting (150 lux), and obtained the same result as that measured by ECG. The measurement accuracy changed as the measurement distance and illumination changed. We set up an illuminance control group and a distance control group, and obtained the relationship between these two variables and the measurement accuracy. We let the environmental lighting switch between low and high illuminance levels. Our analysis of the filtered data verified that our proposed method has strong illumination robustness.

The proposed method offers a non-invasive and non-contact approach to measure heart rate using an ordinary camera. Its ability to avoid the discomfort of traditional measurement methods to the user and ease of operation make it suitable for daily heart rate monitoring. When used with a home network surveillance camera, this method allows real-time observation of the user’s heart rate and calculate the possibility of the user suffering from any CVD. When such possibility reaches a certain value, the user will be reminded to go to the hospital for a medical check. This method has important implications for the prevention of CVDs and is expected to be widely used in nursing homes, workplaces, dormitories, and many other places.

## ACKNOWLEDGMENT

This work was supported by the National Natural Science Foundation of China (Grant No. 61873307 and 61503322), the Scientific Research Project of Colleges and universities in Hebei Province (Grant No. ZD2019305), the Administration of Central Funds Guiding the Local Science and Technology Development (Grant No. 206Z1702G), the Fundamental Research Funds for the Central universities (Grant No. N2023015), the Science and Technology Planning Project of Qin-huangdao (Grant No. 201901B013), and Hebei Natural Science Foundation (Grant No. F2020501040).

---

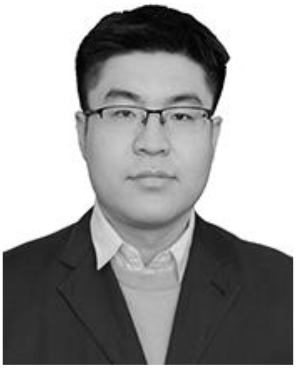

**YIMING YANG** is currently a B.E. student at School of Control Engineering, Northeastern University at Qinhuangdao, China. His current research interests are in the area of deep learning, pattern recognition and computer vision.

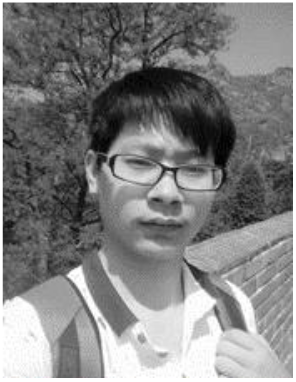

**CHAO LIAN** received the B.E. degree from Liaoning Technical University, in 2017. He is currently pursuing the M.S. degree with the School of Control Engineering, Northeastern University at Qinhuangdao, China. His current research interests are in the areas of wearable cyber physical devices, inertial measurement unit, motion analysis, machine learning, and artificial intelligence.

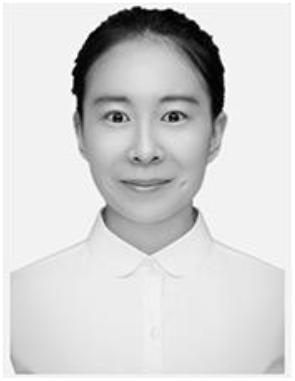

**HONGYU ZHANG** received her B.S. degree in automation from Shenyang Aerospace University, and received the Master degree in control engineering from Yanshan University. She is a research assistant in Northeastern University at Qinhuangdao currently. Her research interests include artificial intelligence, data analysis and electrochemistry

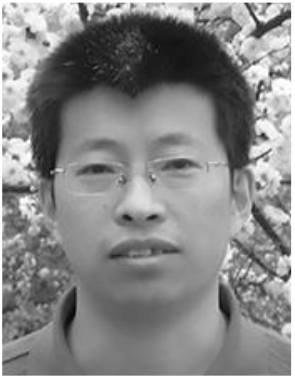

**LIANJIANG LI** received his B.S. degree in physical electronics from the Heilongjiang University, M.S. degree in physical electronics from Harbin Institute of Technology, and Ph.D. degree in physical electronics from Harbin Institute of Technology in 2011. He is currently an Lecturer with the Northeastern University at Qinhuangdao, Qinhuangdao, China. His research interests include Infrared Polarization Imaging, machine learning, infrared simulation, and big data analyses; his plying these technologies to Target recognition and tracking analyses.

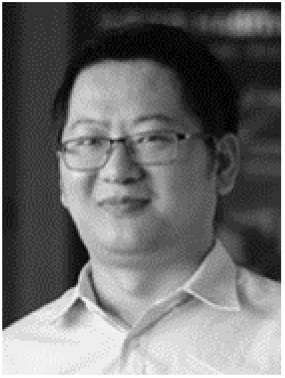

**YULIANG ZHAO** received his B.S. degree in mechanical engineering from the Hubei University of Automotive Technology, M.S. degree in mechanical engineering from Northeastern University, and Ph.D. degree in mechanical and biomedical engineering from the City University of Hong Kong in 2016. He is currently an Assistant Professor with the Northeastern University at Qinhuangdao, Qinhuangdao, China. His research interests include intelligent sensors, machine learning, motion analytics, and big data analyses; his recent work involves applying these technologies to sports and biomechanical analyses.

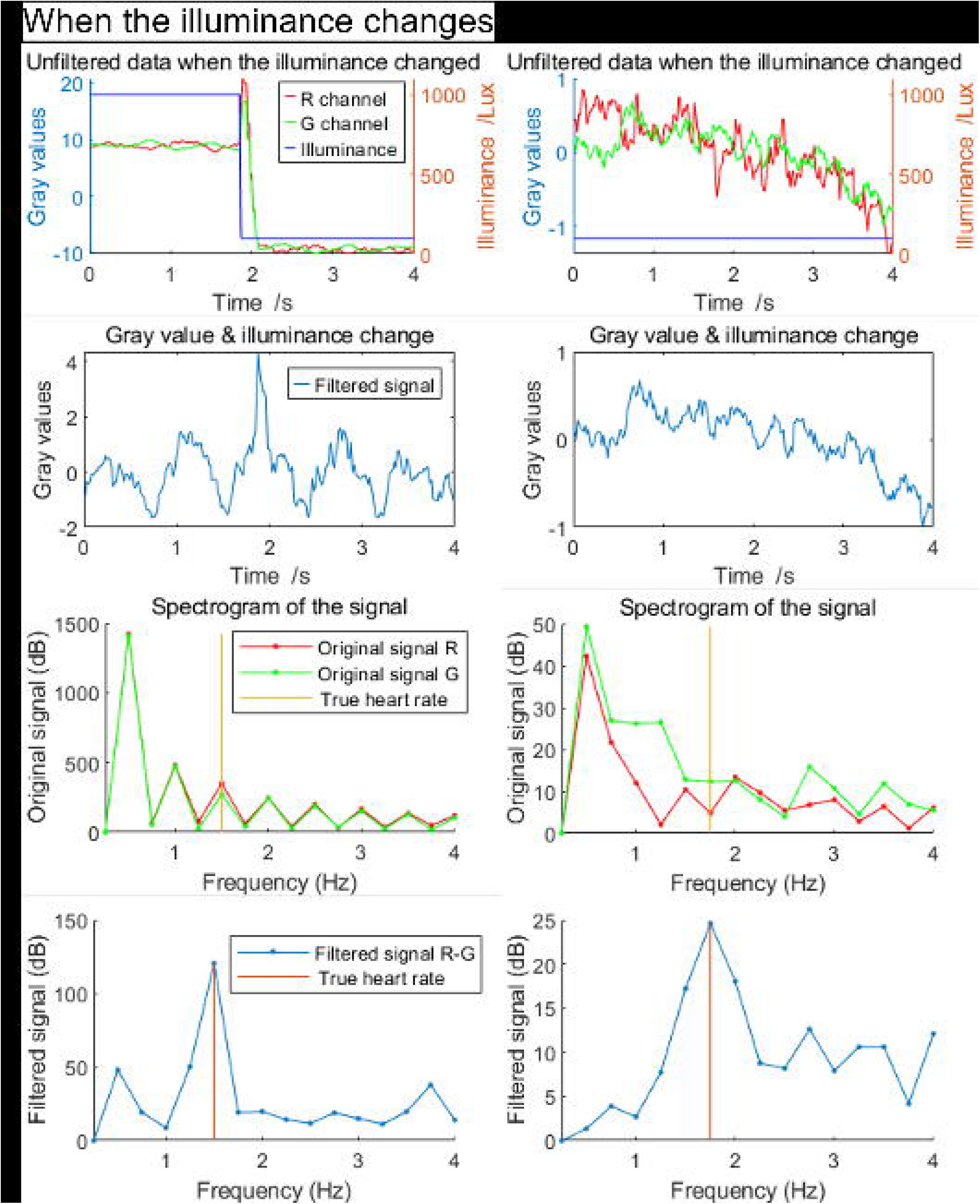

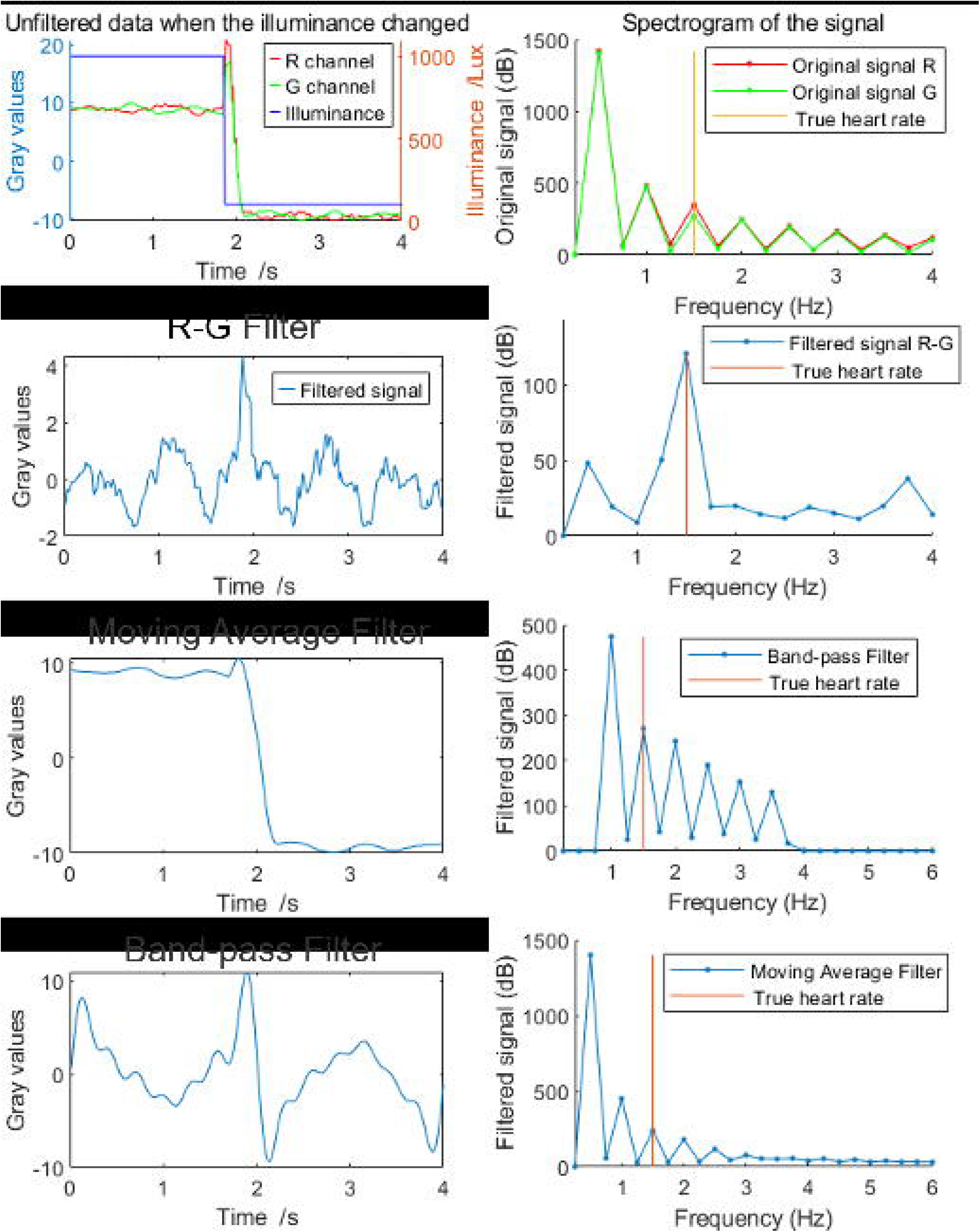

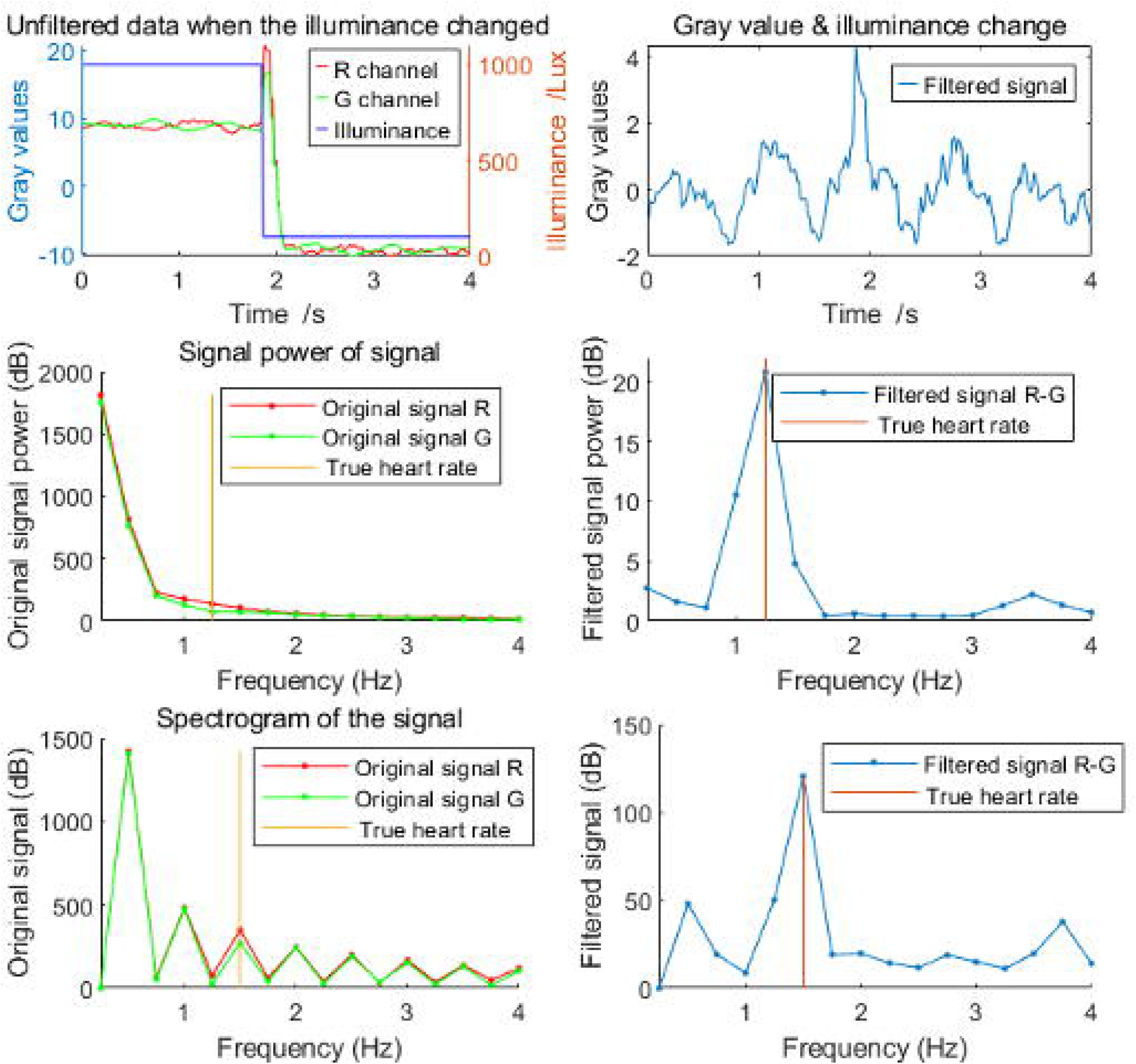

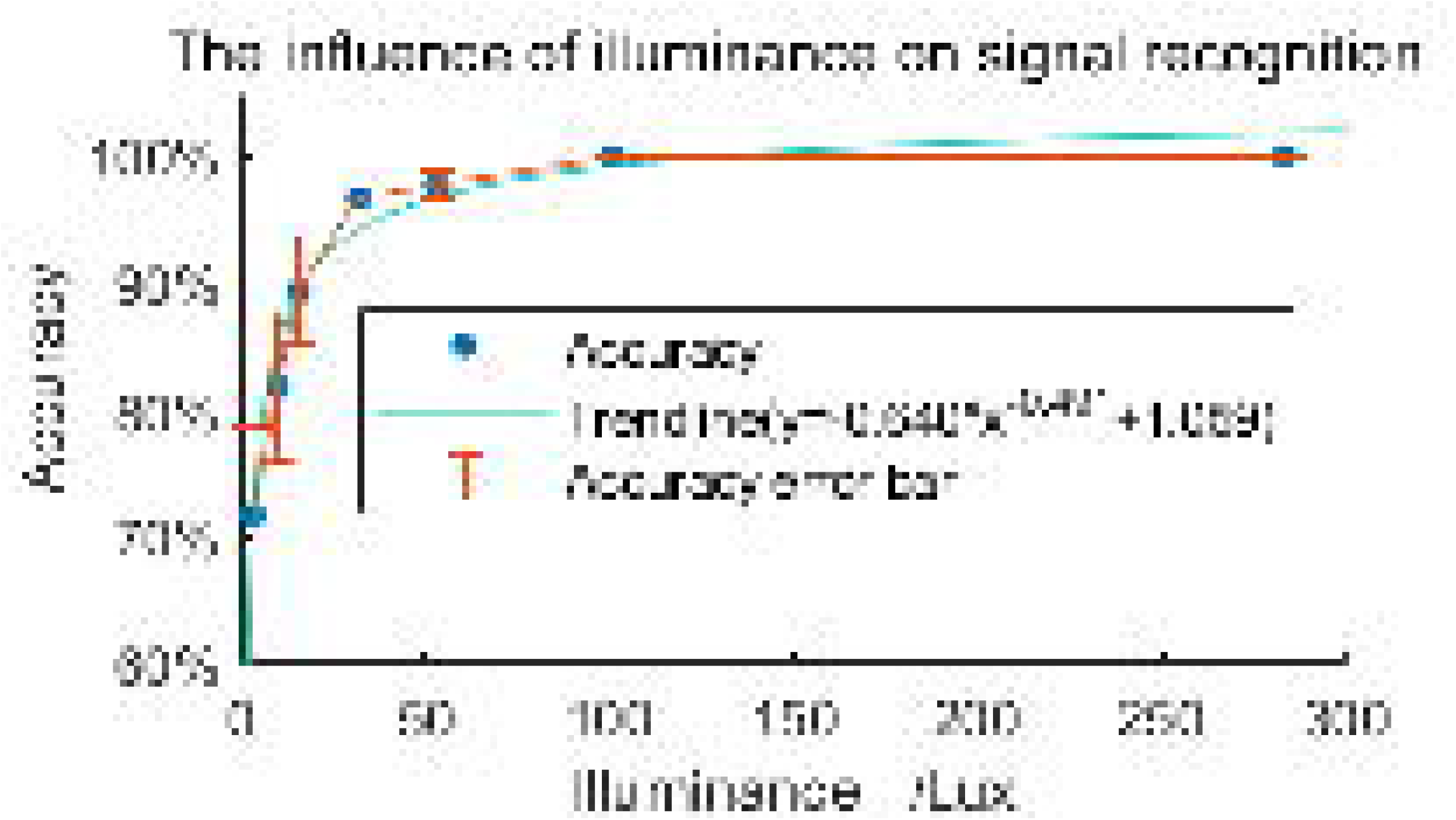

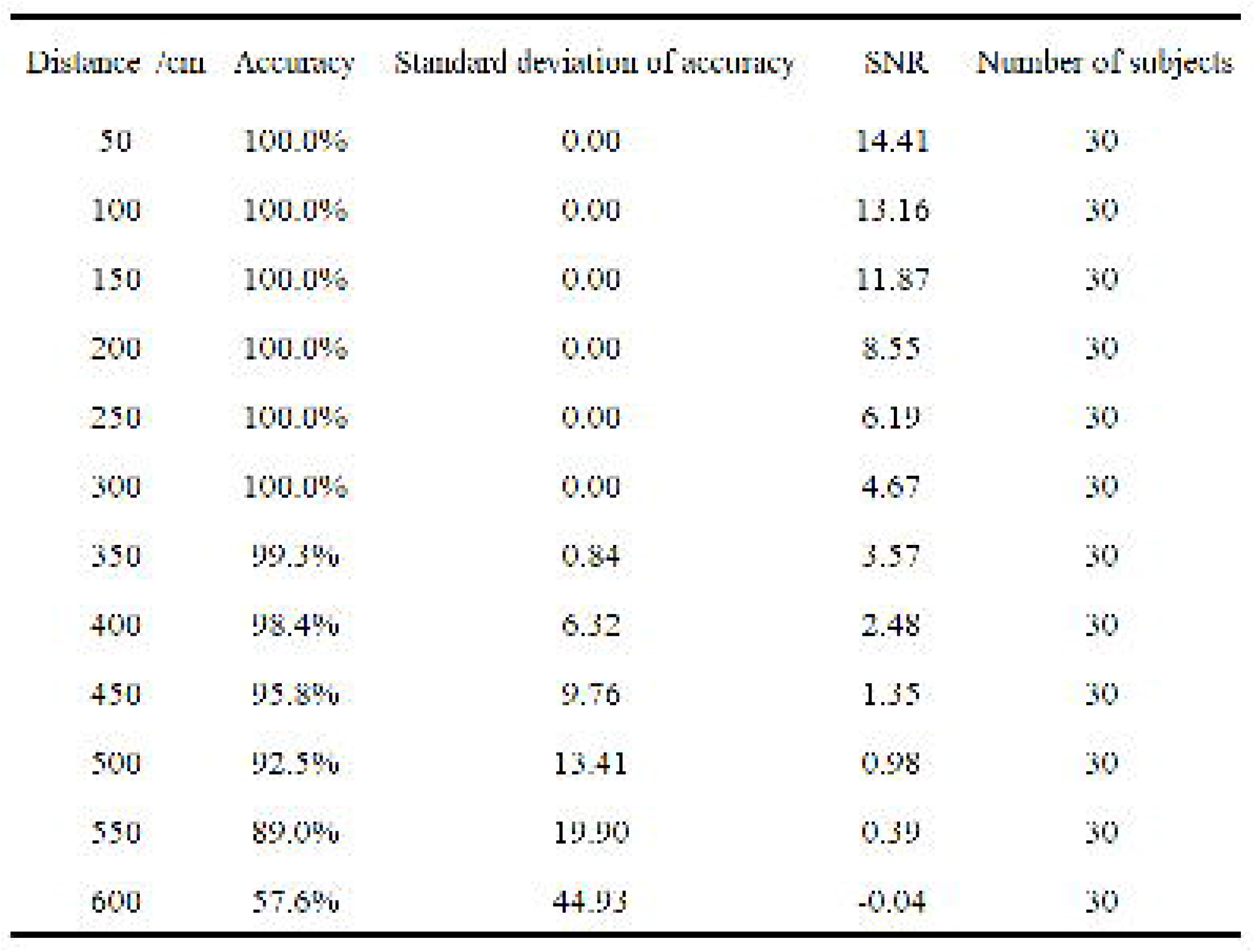

| Distance /cm | Average accuracy (PSD) | Average accuracy (FFT) | Processing time (PSD) /ms | Processing time (FFT) /ms |
| --- | --- | --- | --- | --- |
| 50 | 99.99% | 99.99% | 0.3587 | 0.0157 |
| 100 | 99.99% | 99.99% | 0.3828 | 0.0160 |
| 150 | 99.99% | 99.99% | 0.4478 | 0.0172 |
| 200 | 99.99% | 99.99% | 0.3654 | 0.0157 |
| 400 | 98.41% | 98.41% | 0.3817 | 0.0150 |
| 450 | 95.84% | 95.84% | 0.3817 | 0.0150 |
| 500 | 92.54% | 92.54% | 0.4116 | 0.0134 |
| 550 | 88.98% | 88.98% | 0.3771 | 0.0151 |
| 600 | 57.64% | 57.64% | 0.4426 | 0.0160 |

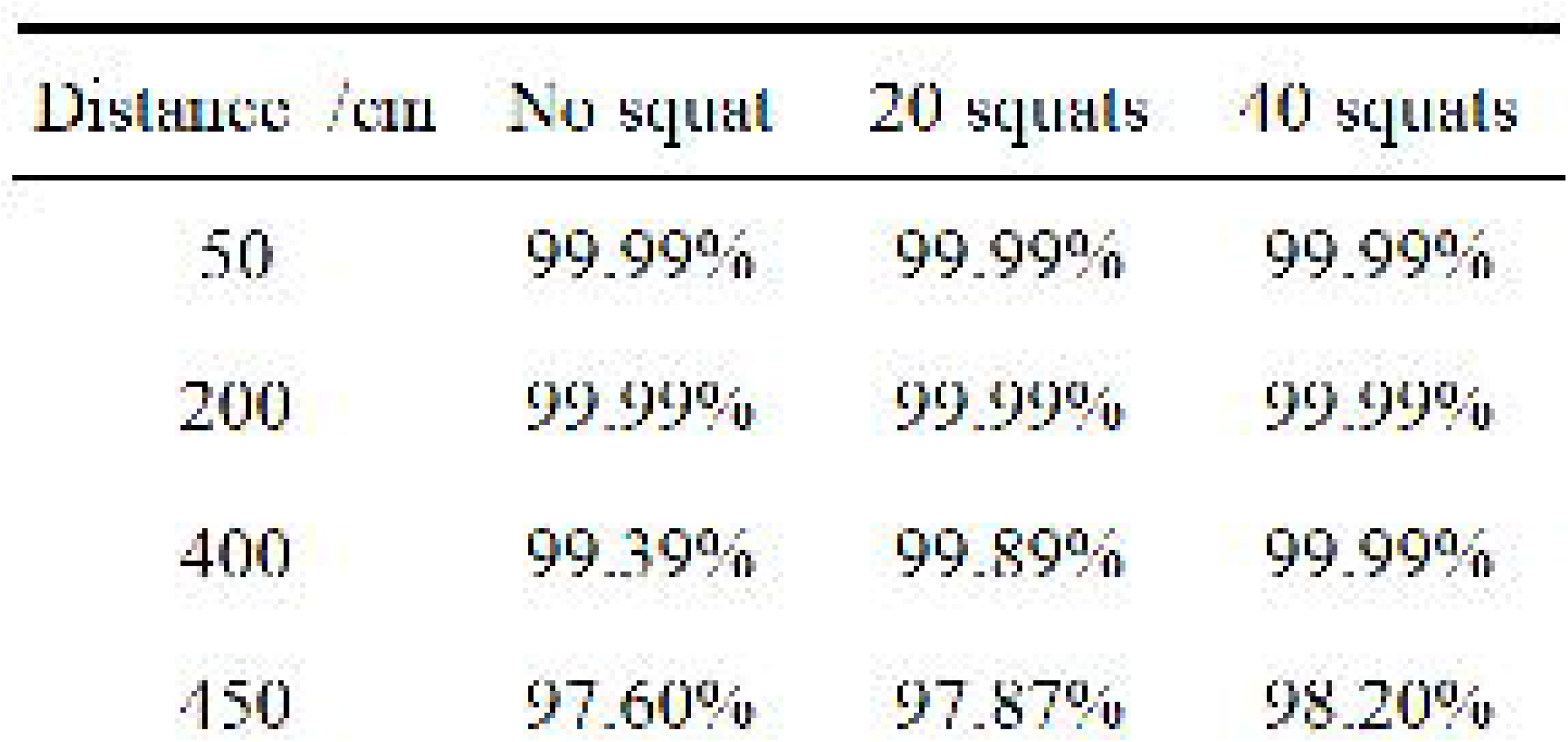

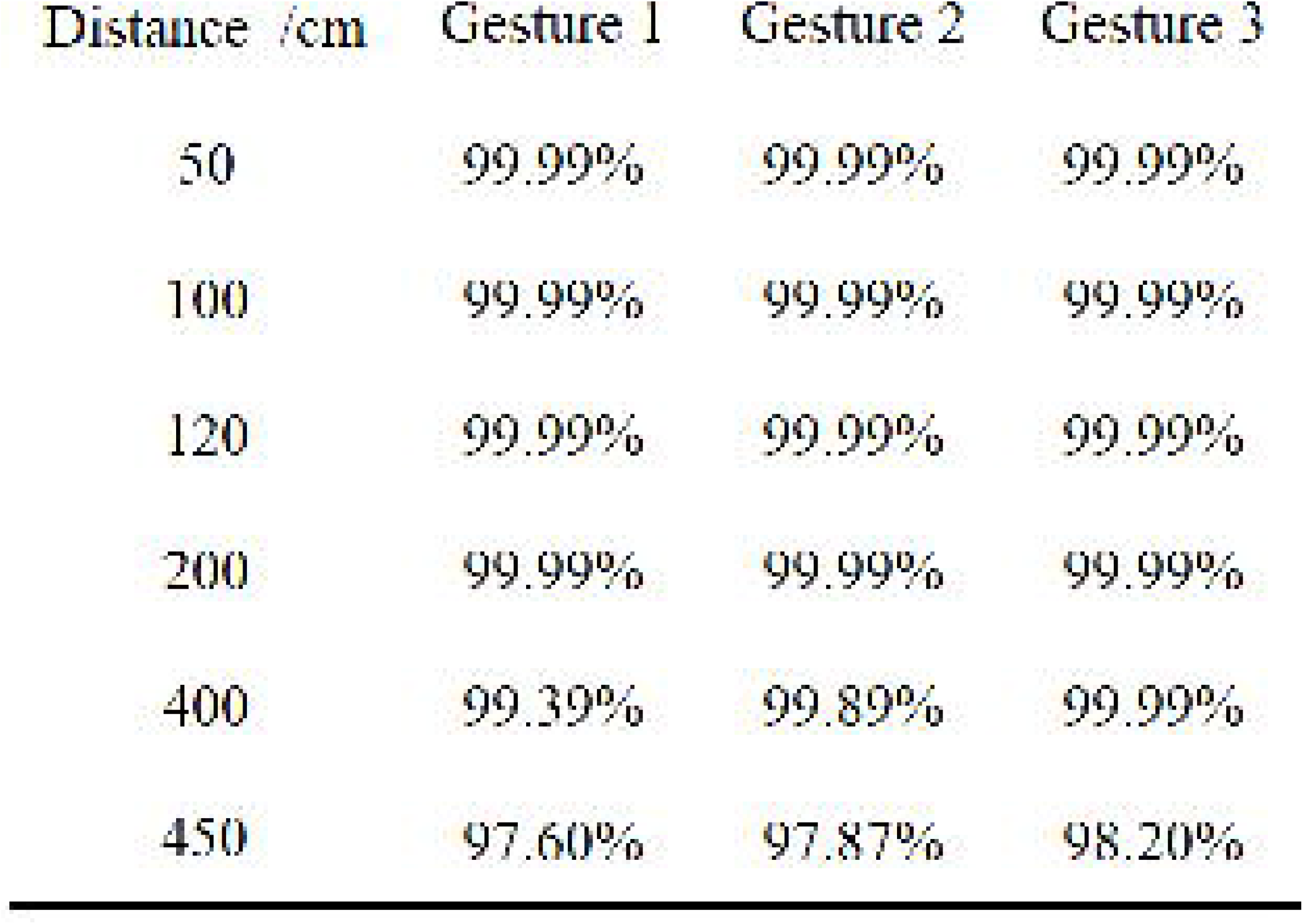

## REFERENCES

[1] S. Petersen, V. Peto, M. Rayner, J. Leal, R. Luengo-Fernandez, and A. Gray, “European Cardiovascular Statistics: 2005 edition,” pp. 1–25, 2005.

[2] M. Nichols, N. Townsend, and M. Rayner, “European cardiovascular disease statistics,”Eur. Hear. Netw. Eur. Soc. Cardiol., pp. 10–123, 2012.

[3] E. Wilkins, L. Wilson, K. Wickramasinghe, and P. Bhatnagar, “European Cardiovascular Disease Statistics 2017 edition,”Eur. Hear. Netw., pp. 94–100, 2017.

[4] https://www.who.int/news-room/fact-sheets/detail/cardiovascular-diseases-(cvds).

[5] R. M. Carney, K. E. Freedland, P. K. Stein, J. A. Skala, P. Hoffman, and A. S. Jaffe, “Change in heart rate and heart rate variability during treatment for depression in patients with coronary heart disease,”Psychosom. Med., vol. 62, no. 5, pp. 639–647, 2000.

[6] G. Mancia et al., “Blood pressure and heart rate variabilities in normotensive and hypertensive human beings,”Circ. Res., vol. 53, no. 1, pp. 96–104, 1983.

[7] M. Böhm et al., “Heart rate as a risk factor in chronic heart failure (SHIFT): The association between heart rate and outcomes in a randomised placebo-controlled trial,”Lancet, vol. 376, no. 9744, pp. 886–894, 2010.

[8] C. B. Taylor, “Depression, heart rate related variables and cardiovascular disease,”Int. J. Psychophysiol., vol. 78, no. 1, pp. 80–88, 2010.

[9] A. Cultur et al., “Covariance structure analysis of health-related indicators in elderly people at home with a focus on subjective feeling of health,”Geographie, vol. 18, no. April 2007, p. 2014, 2009.

[10] E. F. Greneker, “Radar sensing of heartbeat and respiration at a distance with applications of the technology,” in IEE Conference Publication, 1997, no. 449, pp. 150–154.

[11] F. Wurtenberger, T. Haist, C. Reichert, A. Faulhaber, T. Boettcher, and A. Herkommer, “Optimum Wavelengths in the near Infrared for Imaging Photoplethysmography,”IEEE Trans. Biomed. Eng., vol. 66, no. 10, pp. 2855–2860, 2019.

[12] M. Lewandowska, J. Rumiński, T. Kocejko, and J. Nowak, “Measuring pulse rate with a webcam - A non-contact method for evaluating cardiac activity,”2011 Fed. Conf. Comput. Sci. Inf. Syst. FedCSIS 2011, pp. 405–410, 2011.

[13] M. Z. Poh, D. J. McDuff, and R. W. Picard, “Advancements in noncontact, multiparameter physiological measurements using a webcam,”IEEE Trans. Biomed. Eng., vol. 58, no. 1, pp. 7–11, 2011.

[14] D. Djeldjli, F. Bousefsaf, C. Maaoui, and F. Bereksi-Reguig, “Imaging Photoplethysmography: Signal Waveform Analysis,”Proc. 2019 10th IEEE Int. Conf. Intell. Data Acquis. Adv. Comput. Syst. Technol. Appl. IDAACS 2019, vol. 2, pp. 830–834, 2019.

[15] R. H. Goudarzi, S. Somayyeh Mousavi, and M. Charmi, “Using imaging Photoplethysmography (iPPG) Signal for Blood Pressure Estimation,”Iran. Conf. Mach. Vis. Image Process. MVIP, vol. 2020-Febru, pp. 14–19, 2020.

[16] W. Verkruysse, L. O. Svaasand, and J. S. Nelson, “Remote plethysmographic imaging using ambient light,”Opt. Express, vol. 16, no. 26, p. 21434, 2008.

[17] A. Yunitsyna, “Universal Space in Dwelling-the Room for All Living Needs,” no. 131, pp. 8–10, 2014.

[18] S. Zaunseder, A. Trumpp, D. Wedekind, and H. Malberg, “Cardiovascular assessment by imaging photoplethysmography-a review,”Biomed. Tech., vol. 63, no. 5, pp. 529–535, 2018.

[19] A. Özmen, G. W. Weber, I. Batmaz, and E. Kropat, “RCMARS: Robustification of CMARS with different scenarios under polyhedral uncertainty set,”Commun. Nonlinear Sci. Numer. Simul., vol. 16, no. 12, pp. 4780–4787, 2011.

[20] N. Onak, Y. Serinagaoglu Dogrusoz, and G. W. Weber, “Effect of the geometric inaccuracy in multivariate adaptive regression spline-based inverse ECG solution approach,”Comput. Cardiol. (2010)., vol. 44, pp. 1–4, 2017.

[21] T. P. Sacramento, I. M. B. Souza, P. V. O. Vitorino, and T. M. G. A. Barbosa, “A real-time software to the acquisition of heart rate and photoplethysmography signal using two region of interest simultaneously via webcam,”2017 IEEE MIT Undergrad. Res. Technol. Conf. URTC 2017, vol. 2018-Janua, pp. 1–4, 2018.

[22] D. Shao, “Monitoring Physiological Signals Using Camera,”ProQuest Diss. Theses, no. December, p. 108, 2016.

[23] W. G. Zijlstra, A. Buursma, and W. P. Meeuwsen-van der Roest, “Absorption spectra of human fetal and adult oxyhemoglobin, de-oxyhemoglobin, carboxyhemoglobin, and methemoglobin,”Clin. Chem., vol. 37, no. 9, pp. 1633–1638, 1991.

[24] S. Sanyal and K. K. Nundy, “Algorithms for Monitoring Heart Rate and Respiratory Rate From the Video of a User’s Face,”IEEE J. Transl. Eng. Heal. Med., vol. 6, no. February, pp. 1–11, 2018.

[25] G. De Haan and V. Jeanne, “Robust pulse rate from chrominance-based rPPG,”IEEE Trans. Biomed. Eng., vol. 60, no. 10, pp. 2878–2886, 2013.

[26] M. Huelsbusch, ‘Ein bildgestuetstes, funktionelles Verfahren zur optoelektronischer Erfassung der Hautperfusion,’ Ph.D. dissertation, Fakultaet fuer Elektrotechnik un Informationstechnik, RWTH Aachen Univ., Aachen, Germany, p. 70, Jan. 28, 2008.

[27] Y. C. Lin and Y. H. Lin, “A study of color illumination effect on the SNR of rPPG signals,”Proc. Annu. Int. Conf. IEEE Eng. Med. Biol. Soc. EMBS, pp. 4301–4304, 2017.

[28] K. Otsuka et al., ‘Circadian period of human blood pressure and heart rate in clinical health under ordinary conditions,’ [1989] Proceedings. Second Annual IEEE Symposium on Computerbased Medical Systems, Minneapolis, MN, uSA, 1989, pp. 206–213.

[29] S. Di Pilla, “U.S. Standards and Guidelines,”Slip, Trip, Fall Prev., pp. 161–194, 2009.

[30] K. Niizeki and Y. Miyamoto, “Phase-dependent heartbeat modulation by muscle contractions during dynamic handgrip in humans,”Am. J. Physiol. - Hear. Circ. Physiol., vol. 276, no. 4 45-4, pp. 1331–1338, 1999.

[31] G. Giuriato et al., “Timed synchronization of muscle contraction to heartbeat enhances muscle hyperemia,”J. Appl. Physiol., vol. 128, no. 4, pp. 805–812, 2020.

[32] A. Pears, “Strategic Study of Household Energy and Greenhouse Issues a Report for Environment Australia,”Solutions, no. June, pp. 1–110, 1998.

[33] Schlyter, P. (2010). Radiometry and photometry in astronomy. Retrieved October.

